# Early detection of biomarkers for circulating tumor cells in Bone marrow and Peripheral blood in a fast-progressing gastric cancer model

**DOI:** 10.1101/2020.01.29.925727

**Authors:** Prerna Bali, Ivonne Lozano-Pope, Collin Pachow, Marygorret Obonyo

## Abstract

*Helicobacter pylori* poses one of the greatest risks for development of gastric cancer. We previously established a crucial role for myeloid differentiation primary response 88 (MyD88) in the regulation of *Helicobacter*-induced gastric cancer. Mice deficient in *Myd88* rapidly progressed to neoplasia when infected with *H. felis*, a close relative of *H. pylori*. For this study we examined circulating tumor cells (CTCs) by measuring expression of cytokeratins, epithelial to mesenchymal transition (EMT) and cancer stem cell (CSC) markers in in the bone marrow and peripheral blood of gastric cancer models we termed fast (*Myd88*^-/-^)- and slow (WT)-“progressors”. We detected cytokeratins CK8/18 as early as 3 months post infection in the fast “progressors”. In contrast, cytokeratins were not detected in slow “progressor” gastric cancer model even after 7 months post infection. Expression of MUC1 was observed in both bone marrow and peripheral blood at different time points suggesting its role in gastric cancer metastasis. Snail, Twist and ZEB were expressed at different levels in bone marrow and peripheral blood. Expression of these EMT markers suggests manifestation of cancer metastasis in the early stages of disease development. Lgr5, CD44 and CD133 were the most prominent CSC markers detected. Detection of CSC and EMT markers along with cytokeratins does reinforce their use as biomarkers for gastric cancer metastasis. This early detection of markers suggests that CTCs leave primary site even before cancer is well established. Thus, cytokeratins, EMT, and CSCs could be used as biomarkers to detect aggressive forms of gastric cancers. This information will be important in stratifying patients for treatment before the onset of severe disease characteristics.

## Introduction

Gastric cancer is the third leading cause of cancer mortality worldwide [1]. *Helicobacter pylori* (*H. pylori*) infection is the strongest risk factor for the development of gastric cancer leading to the recognition of this bacterium by the World Health Organization (WHO) as class 1 carcinogen [2–4]. *H. pylori* infections affect up to 80% of the population in certain parts of the globe [3, 5]. Deaths from gastric cancer like most common cancers are a result of metastasis [6]. Moreover, there are no effective predictors for identifying recurrence and metastasis in gastric cancer. Consequently, determination of factors that indicate existence of metastasis is critical for therapeutic interventions with the goal of improving disease outcome. Cancer metastasis involves tumor cells referred to as circulating tumor cells (CTCs), leaving the original cancerous site by migrating to distant sites and these can be found in peripheral blood and bone marrow [7–10]. In the bone marrow, these CTCs are referred to as disseminated cancer cells (DTCs). CTCs have been used as biomarkers of metastasis in many cancer types [9, 10] and their presence is associated with poor prognosis [11] [6, 12–14]. In the bone marrow, evidence of cancer cells at the time of surgical intervention has been associated with metastasis [9, 5]. While studies associating CTCs and DTCs with cancer metastasis have been very well studied for breast and lung cancer [6], not very much has been described for gastric cancer. Indeed, the CellSearch System (Veridex, NJ) was approved by the US Food and Drug Administration for the detection of CTCs in patients with breast, prostate, and colorectal cancer [10, 16–18], and its use for detection of CTCs in gastric cancer continues to be controversial [19]. This has led to a lack of enthusiasm in studies to detect CTCs in gastric cancer patients and consequently their routine usage in gastric cancer management. One of the most common ways to detect CTCs in solid tumors is with the use of surface markers such as cytokeratins (CK) and Mucin-1 (MUC1). Cytokeratins in general have been extensively studied in epithelial cancers such as breast cancer [20] specifically cytokeratins such as CK-8, CK-18, and CK-19 [21]. These markers are of particular interest, due to their abundant expression in epithelial cells and relatively low or no expression in mesenchymal cells [1, 2]. Recently, other markers such as epithelial-to-mesenchymal transition (EMT) and cancer stem cells (CSCs) have been shown to be major components of CTCs due to their association with cancer progression [23–28].

EMT, which depicts change in epithelial cells to a malignant phenotype [29] is considered a crucial step in cancer progression [30]. This process disrupts crucial activities such as cell-cell adhesion, cell polarity [31], and extra cellular matrix degradation [32]. There are many inducers of EMT, most notably factors such as cytokines, innate and adaptive immune responses, and growth factors secreted by tumor microenvironment among others [3, 4]. This EMT process is tightly regulated by transcription factors such as Snail, Twist, and ZEB. Snail and Twist have previously been shown to be over expressed in *H. pylori-infected* patients [35]. While patients with early stages of cancer do not exhibit EMT phenotypes, gastric cancer cell motility and metastasis was observed in the advanced stages of gastric cancer which was implicated in the EMT process [36]. While the clinical significance of EMT in other cancers has been confirmed [29, 37, 38], in gastric cancer, although the expression of EMT-related proteins has been studied, their significance is still questionable [39, 40]. CSCs, which are also suggested to be components of CTCs are believed to contribute to many of the aggressive cancer characteristics such as metastasis, tumor invasion, chemotherapy resistance and relapse [41].

Knowledge of micrometastatic cells including when they arise and their detection is critical since their dissociation from the primary tumor microenvironment and transportation to distant sites and finally colonization is what ultimately leads to death. These cells are therefore important for the detection of metastasis or disease recurrence. Detection of CTCs is therefore crucial in identifying patients that are likely to relapse or develop metastases and can subsequently be targeted for suppression of metastasis. Our previous studies have shown that absence of myeloid differentiation primary response gene 88 (MyD88) leads to development of an aggressive form of *Helicobacter*-induced gastric cancer, resulting in gastric cancer mouse models we termed slow (wild type) and fast (*Myd88*^-/-^) “progressors” [2, 3]. In the present study, we used these gastric cancer models to evaluate kinetics of CTCs and DTCs over a span of 6 months. We detected them using surface markers, cytokeretins and mucins; EMTs and CSC markers, in the bone marrow and peripheral blood by employing immunocytochemistry (ICC), and/or quantitative real-time polymerase chain reaction (qRT-PCR). Thus, data from this study indicate that early detection of metastasis in aggressive gastric cancers maybe useful for patients by providing proper prognosis and treatment.

## Material and Methods

### Animals

Mice aged six to ten weeks old were used in this study. Wild type (WT) and *Myd88*^-/-^ mice in C57BL/6J background were purchased from the Jackson Laboratory (Bar Harbor, ME). In addition, some *Myd88*^-/-^ mice were bred in house. All mice were housed together for the duration of the study and prior to *H. felis* infection. All animal procedures were approved by the Institutional Animal Care and Use Committee at the University of California, San Diego. All procedures were performed using accepted veterinary standards.

### Bacterial Strains and Growth Conditions

*H. felis* strain CS1 (ATCC 49179) was used for mouse infections. This strain was originally purchased from American Type Culture Collection (Manassas, VA). *H. felis* were maintained on both solid and liquid medium. The solid medium was composed of Columbia agar (Becton Dickson, MD) supplemented with laked blood (5%, Hardy Diagnostics, CA) and Amphotericin B (1%; Mediatech, VA). The liquid medium was composed of Brain Heart Infusion (BHI; Becton Dickson, MD) supplemented with 10%, heat inactivated fetal bovine serum (FBS).

Bacteria were grown at 37°C under microaerophilic conditions (5% O_2_, 10% CO_2_, 85% N_2_) as described in our previous studies [2, 4]. Bacteria maintained on solid media were passaged every 2-3 days. Before infections of mice, *H. felis* were grown in liquid media for 48 hrs. Spiral bacteria were counted using a Petroff-Hausser chamber.

### Mouse Infections

WT and *Myd88*^-/-^ mice were infected with *H. felis* grown in BHI. A total of 10^9^ organisms in 300μl were administered to each mouse by oral gavage every other day for a total of 3 inoculations as described in our previous studies [44] [42]. Control mice received 300 μl of BHI. *Myd88*^-/-^ mice (2 or 3 mice) were euthanized every month up to 6 months post infection. WT mice (2 or 3 mice) were euthanized at 5, 6, and 7 months post infection. Bone marrow and peripheral blood was aseptically removed and processed for experimental analysis.

### Bone marrow Isolation

Following euthanasia, femurs and tibias were aseptically removed from mice taking care to remove any muscle on or near the bones as described in our previous studies [5, 6]. Bone marrow cells were flushed using a 22-gauge needle and phosphate buffered saline (PBS) by cutting the ends of the bones with sharp scissors. Cells were collected for downstream applications.

### Immunocytochemistry

Samples collected from bone marrow and peripheral blood were deposited onto lysine coated slides using StatSpin CytoFuge (Beckman Coulter, Indianapolis, IN). Briefly, cell samples were loaded onto cell concentrators with lysine coated slides. The concentrators with sample were then placed into the cytofuge and spun at 55 x g for 4 min. Once cell samples were placed on slides, cells were fixed with 4% paraformaldehyde for 10 min. Cells were then incubated with 1% bovine serum albumin (BSA) in 0.1% PBS supplemented with Tween 20 (PBST) for 30 min. Cells were immunostained with antibodies specific for CK8/18 (EP1628BY, 1:2000, Abcam (Cambridge, MA)) and CD117 (c-Kit, 2B, 1:200, Santa Cruz Biotechnologies (Santa Cruz, CA)). After washes, cells were incubated with anti-rabbit secondary antibody (1:1000) with fluorochrome for 1 hour in the dark and the samples were then mounted with Fluoroshield mounting medium with DAPI (4’,6-diamidino-2-phenylindole, Abcam, Cambridge, MA). Slides were imaged using the Keyence BZX-700 Fluorescent Microscope (UCSD Microscopy Core).

### Isolation of RNA and cDNA synthesis

RNA was isolated from bone marrow and peripheral blood samples using Direct-zol RNA mini kit (Zymo Research, Irvine CA) according to the manufacturer’s instructions. Briefly, a total of TRI Reagent was added to bone marrow or blood plasma in a volume of 3:1. The samples were vortexed vigorously followed by RNA purification. The samples were passed through a collection column and washed with the accompanying buffers. The resulting RNA solution was passed through a filter cartridge and RNA eluted using nuclease-free water. RNA quality was determined using a Nanodrop system (Thermo Fisher, Waltham MA) by reading absorbance levels at 260 nm. 2μg of RNA per sample was reverse transcribed into cDNA using High Capacity cDNA Reverse Transcription Kit (Thermo Fisher, Waltham MA).

### Quantitative real time PCR

Quantitative real time PCR (qRT-PCR) was performed as described in our previous studies [44, 46, 47]. We determined the expression of select genes including, CD44, SOX9, Prominin-1 (CD133), SOX2, OCT4, NANOG, Lgr5, CK-18, CK-19, MUC1, Snail, Twist, and ZEB. Briefly, 2μl of cDNA was used per well in a total of 10μl reaction mix for amplification using

Step One Real Time PCR (Applied Biosystems, Carlsbad CA). The amplification conditions consisted of an initial cycle of 95°C for 5 minutes followed by 40 cycles of amplification with denaturation as follows: 95°C for 15 sec, 60°C for 20 sec, 72°C for 40 sec. Gene expression levels were normalized to Glyceraldehyde 3-phosphate dehydrogenase (GAPDH). The data collected was analyzed using comparative cycle threshold calculations (ΔΔC_t_, Applied Biosystems) and plotted using GraphPad Prism software. Primers used are listed in Supplementary Table 1.

## Results

### Detection of epithelial markers in bone marrow and peripheral blood in response to *Helicobacter* infection

Epithelial markers CK8/18 were used to detect CTCs and DTCs in peripheral blood and bone marrow of mice in response to infection with *H.felis*, respectively. c-Kit (CD117) was used as a standard surface marker expressed in hematopoietic cells and progenitor cells in the bone marrow [48] [49]. In our fast “progressor” gastric cancer model, bone marrow was analyzed for epithelial markers monthly up to 6 months post infection. Epithelial cell markers were detected in the bone marrow as early as 3 months and their expression levels increased as the disease progressed with maximum expression observed at 6 months post infection (Fig. 1). These markers were not detected at 1 (Fig. 1) or 2 months (Fig. S1). We did not detect CK8/18 in peripheral blood. However, increased expression of CK-18 and CK-19 in both peripheral blood and bone marrow (Fig. 2) was observed as the disease progressed using qRT-PCR. Moreover, Mucin 1(MUC1) expression was observed in peripheral blood both at 4 and 6 months whereas its expression in bone marrow was only observed at 6 months (Fig. 2B). It has been reported that expression levels of CTCs are generally lower in peripheral blood compared to bone marrow [50]. On the other hand, no epithelial markers were detected in the slow “progressor” gastric cancer model *(H.* felis-infected WT mice) at 5, and 6 months post infection (Fig. S2). Therefore, all subsequent experiments were only performed in the fast “progressor” gastric cancer model (*H. felis*-infected *Myd88*^-/-^ mice).

**Figure 1.**
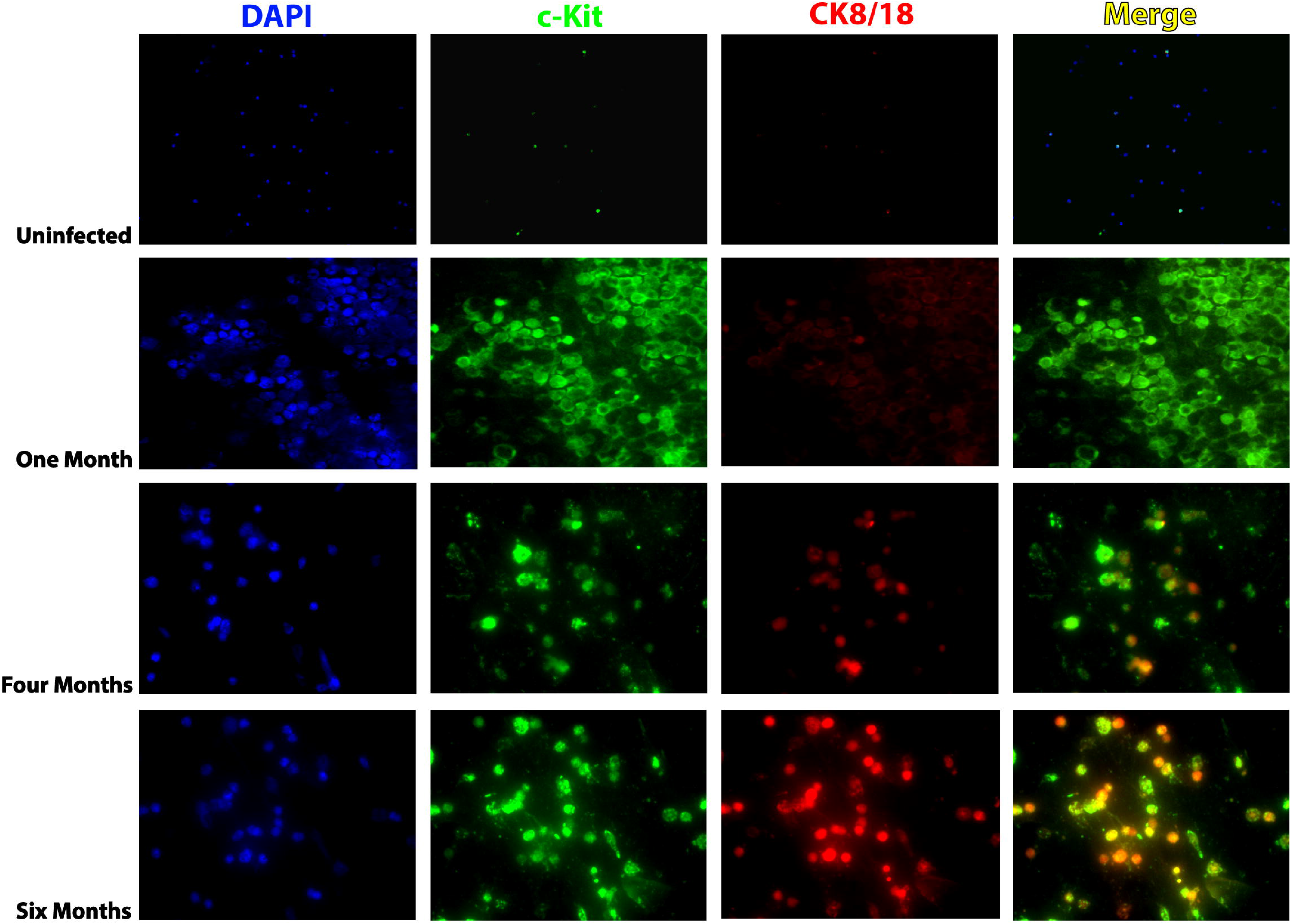
Infection with *Helicobacter felis* induces epithelial marker expression in the fast “ progressor” gastric cancer model *(H.* felis-infected Myd88^-/-^mice). Representative images of immunofluorescent staining of bone marrow for c-Kit (green) and CK-8/18 (red) are shown with DAPI staining of nuclei in blue. Myy88^-/-^ mice were infected with *H. felis*, and mouse bone marrow was checked monthly from 1 to 6 months to determine epithelial marker expression. Images shown are uninfected (a), 1 months post-infection (b), 4 months postinfection (c), and 6 months post-infection (d).

**Figure 2.**
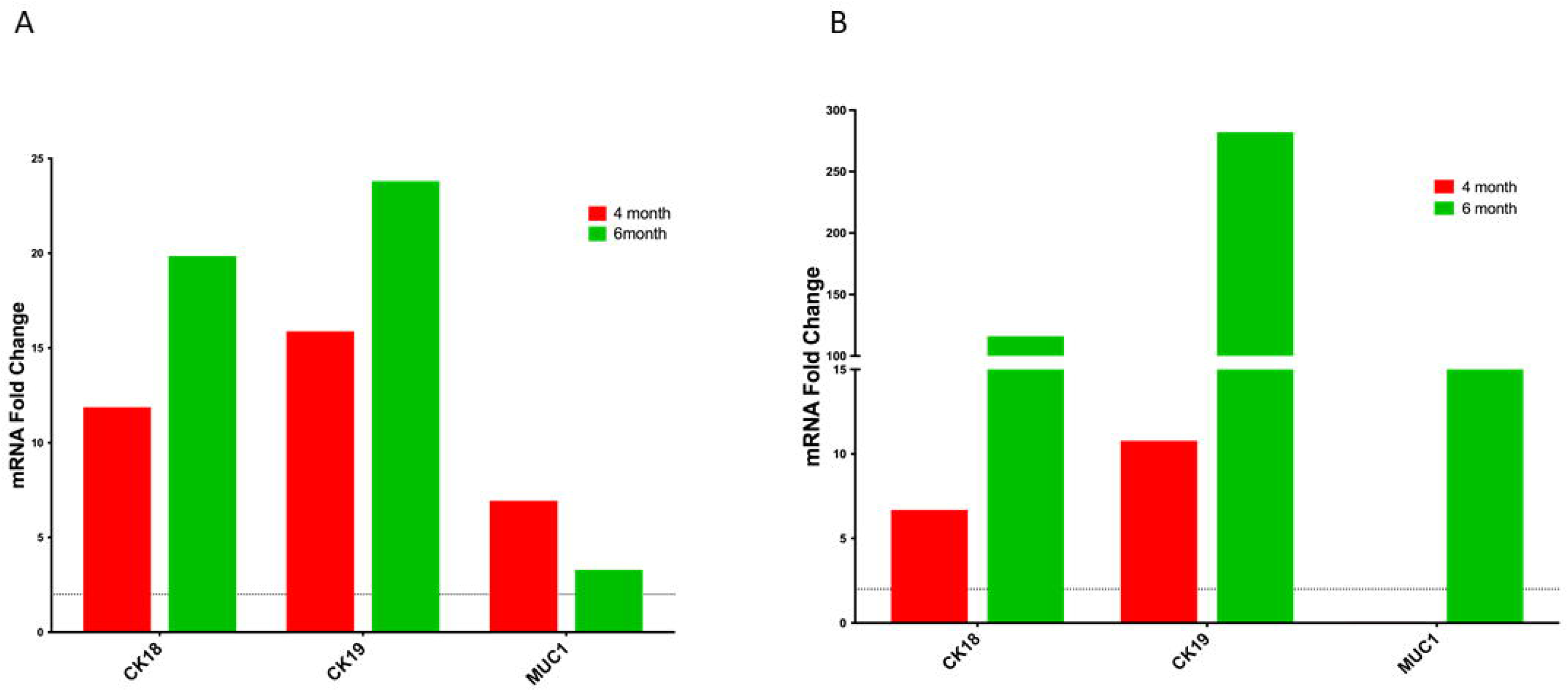
Epithelial markers in bone marrow and peripheral blood are overexpressed. Gene expression of CK-18, CK-19, and MUC1, in peripheral blood (a) and bone marrow (b) from Myd88^-/-^ *H. felis-infected* mice. Data are reported as fold induction vs Myd88^-/-^ uninfected mice.

### Evidence of epithelial transition to a mesenchymal phenotype

We analyzed EMT transcription factors Snail, Twist and ZEB to determine their expression during H.felis-induced disease progression. Increased expression levels of Snail was observed in bone marrow as compared to negligible or below threshold levels at both 4 months and 6 months in peripheral blood (Fig. 3). On the other hand, although expression levels of Twist was observed above threshold levels at both 4 and 6 months post infection, the peak levels were different, they peaked at 6 months in peripheral blood (Fig. 3A) and at 4 months in bone marrow (Fig. 3B). ZEB was expressed in peripheral blood but was undetectable in bone marrow (Fig. 3).

**Figure 3.**
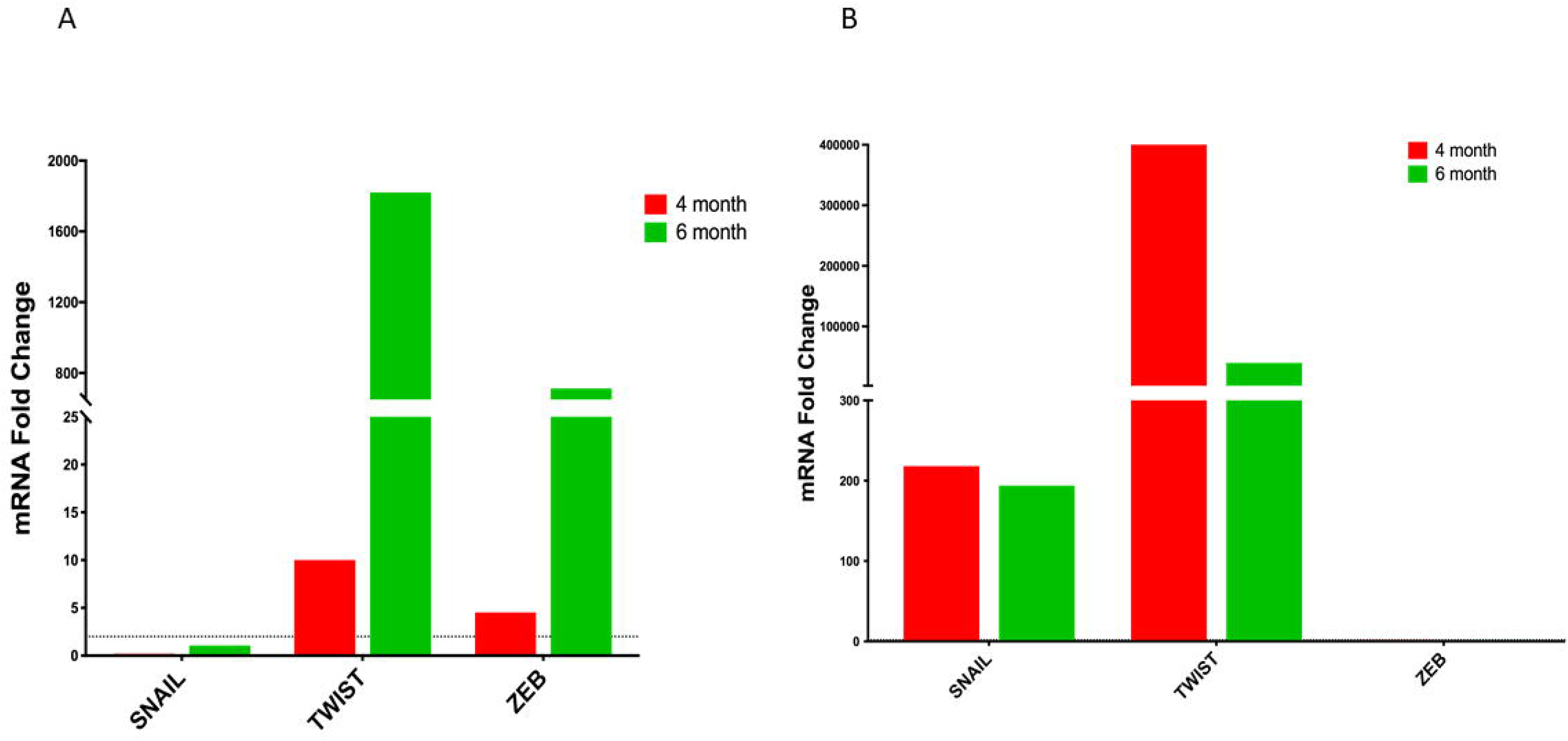
Expression of epithelial-mesenchymal transition (EMT) markers. Snail, Twist, and ZEB levels were assessed in bone marrow (a) and peripheral blood (b) from Myd88^-/-^ *H.* felis-infected mice at 4- and 6-months using RT-PCR. Data are reported as fold induction vs Myd88^-/-^ uninfected mice.

### Expression of cancer stem cells markers in bone marrow and peripheral blood

Lgr5 is the most well-known gastric cancer stem cell marker and has been studied extensively to validate its importance in gastric cancer [51]. In our study expression of Lgr5 increased gradually from 4 to 6 months in peripheral blood and bone marrow with higher expression levels in the bone marrow (Fig. 4). Expression of CD44, which was the first gastric cancer stem cell marker identified [2, 3] peaked at 6 months in the bone marrow. On the other hand, CD133 was expressed at high levels in peripheral blood (Fig. 4B) but was undetectable in the bone marrow (Fig. 4A). Other cancer markers evaluated included, OCT4, NANOG, SOX2, and SOX9 and their expression levels were detected in both bone marrow and peripheral blood (Fig. 4).

**Figure 4.**
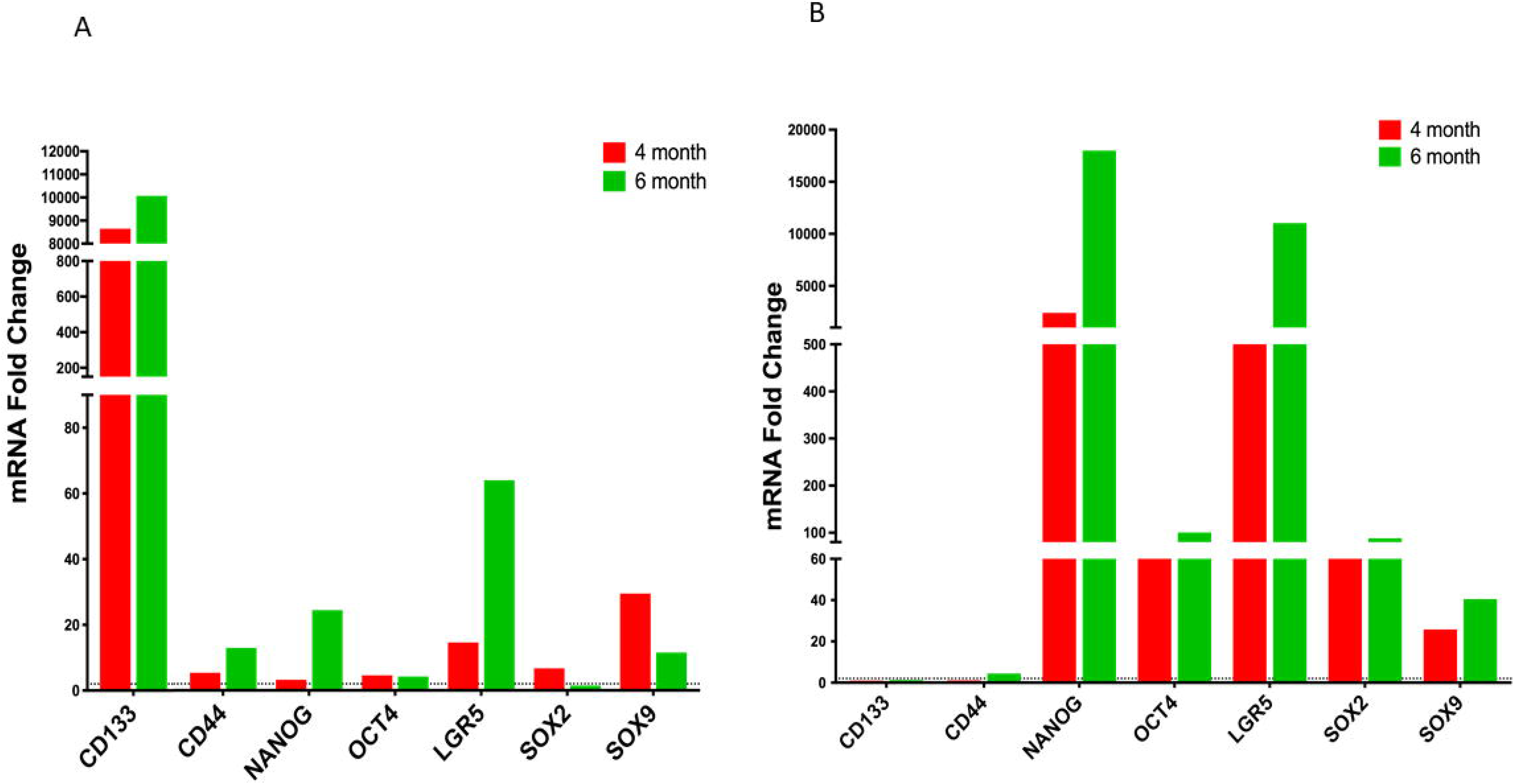
Quantification of cancer stem cell marker levels. CD133, CD44, NANOG, OCT4, LGR5, SOX2, and SOX9 levels were assessed in bone marrow (a) and peripheral blood (b) from Myd88^-/-^ *H. felis-infected* mice at 4- and 6-months using RT-PCR. Data are reported as fold induction vs Myd88^-/-^ uninfected mice.

## Discussion

Examination and diagnostic tools for confirming the presence of gastric cancer are often invasive with endoscopy being the main test used to detect stomach cancer. At times, signs and symptoms are not very distinguishable for many patients and with no protocol in place in countries where incidence of gastric cancer is low, the chance of early stage detection is very minimal. For most patients, gastric cancer is diagnosed in the locally-advanced or late stages because either screening was not performed or the disease was detected only after the development of symptoms. Early detection will help increase patient survival by decreasing the chance for metastatic progression. Indeed, detection of CTCs in peripheral blood and bone marrow in gastric cancer patients has been suggested to be indicative of metastasis [4, 5]. However, clinical significance of CTCs and DTCs as indicators of metastasis has not been appropriately utilized in gastric cancer compared to breast and lung cancer [6]. In our present study we utilized three subset of biomarkers – cytokeretins, EMTs, and CSCs to detect CTCs and DTCs indicative of metastasis using our previously established fast “progressor” gastric cancer model at an early stage [42]. In the fast “progressor” model (*Myd88*^-/-^) CTCs were detectable as early as 3 months compared to our slow “progressor” model (WT type) where they were undetectable even at 6 months. This suggests that in an aggressive form of cancer the transformed cells, CTCs start moving to secondary locations even before the cancer is well established at the primary site. Presence of epithelial gastric surface markers within bone marrow and peripheral blood indicate that not only have tumor-like cells left the microenvironment of the gastric mucosa but have successfully begun infiltrating these areas leading to micro metastatic tumors throughout the body [20]. CK-8, CK-18 and CK-19 have previously been identified as markers whose expression is found in almost all epithelial-based carcinomas [9, 6]. Cytokeratins such as 8 and 18 are found in over 90% of gastric cancer tumors [57] making them reasonable targets to evaluate as positive markers of gastric metastasis. In addition, we detected MUC1 in both the peripheral blood and bone marrow. MUC1 is an oncoprotein found in many adenocarcinomas [59], which under normal conditions is known to protect the gastric epithelium [58] [59] [60, 1]. However, in the presence of *H. pylori*, MUC1 expression has been shown to be considerably decreased [59]. MUC1 is one of the markers used for detecting CTCs and DTCs in epithelial solid cancers [6] and has been linked to cancers such as non-small cell lung cancer [62] as well as DTCs within the bone marrow of breast cancer patients [63]. The role of MUC1 in carcinogenesis has not been well elucidated and especially its role in gastric cancer is contradictory [64–66], but its overexpression has been associated with cancer metastases [67]. Recent studies have suggested regulation of MUC1 by mir-206 inhibits proliferation and migration of gastric cancer cells [68]. Thus, reinforcing the role of MUC1 as a gastric cancer metastases biomarker. Moreover, MUC1 promotes cell proliferation by Wnt signaling pathway and EMT activation through Snail in renal carcinoma [69].

EMT is described as transition of cells from epithelial to a mesenchymal state that is associated with suppression of E-cadherin resulting in an invasive cell phenotype [28–30, 70, 71]. This change in expression is induced by EMT-transcription factors (EMT-TFs), which include Snail, Twist and ZEB. Increased expression of these EMT markers is associated with the transition of the epithelium into a malignant phenotype [29]. As gastric cancer progresses, epithelial cells begin to lose these phenotypic markers and begin to acquire a mesenchymal phenotype [72], which is associated with loss of cell-cell adhesion of epithelial cells as well as changes in cell polarity which eventually allows for easier migration of cells [71]. The concomitant expression of these EMT markers with epithelial markers in our gastric cancer model indicates that these EMT markers may be used as indicators of metastasis in gastric cancer. Recent findings suggest that acquisition of mesenchymal markers in tumors is a poor prognostic cancer factor [28, 73, 74]. Hypoxic conditions in tumors are suggested to trigger mesenchymal stem cell (MSCs) migration [75–78]. The presence of these MSCs in tumor stroma is associated with EMT stimulation. Once stimulated it is indicated that these MSCs may promote cell invasion and spread of tumor cells in systemic circulation [79]. As an example, studies carried out by Yang et al., 2004 [80], in breast cancer have suggested that high levels of continued expression of Twist is essential for metastasis. Our findings also show continued expression of Twist in both peripheral blood and bone marrow, thus, suggesting a role of Twist in metastasis in an aggressive form of gastric cancer. Snail is a strong suppresser of E-cadherin and is closely associated with cancer metastasis and tumor progression via the Wnt pathway [81]. Previous studies in breast cancer have shown that Snail is required for lymph node metastasis [29]. High expression levels of Snail in the bone marrow may indicate DTCs in the bone marrow. ZEB in addition to its function as an EMT inducer, also plays a role in hematopoiesis. ZEB has been associated with acquisition of cancer stem cell (CSC) properties

[82]. Thus, expression of ZEB as well in our study indicates early metastasis in a fast progressing gastric cancer.

In this study we also detected CSCs including Lgr5, CD44, CD133, OCT4, SOX2, SOX9, and NANOG in peripheral blood and bone marrow during disease progression. Cancer stem cells play a vital role in cancer metastasis [3, 1]. For all tumor associated cell markers we detected in our fast progressing gastric cancer model, expression levels were always greater in the bone marrow than in the peripheral blood. This is in line with reports on challenges associated with detection of CTCs in blood due to very low numbers of tumor cells in blood

[83]. Indeed, the levels of CTCs are generally lower in peripheral blood compared to bone marrow [50]. Interestingly, CD133 was highly expressed in peripheral blood and undetectable in the bone marrow. The reason for this differential expression remains to be investigated. CD133 is a known cancer stem cell marker in cancers such as colorectal cancer and liver cancer and for is its role in metastasis in these cancers [84]. Of all these CSC markers, Lgr5 and CD44 are well known targets of the Wnt signaling pathway and have been implicated in cancer invasion and metastasis through their involvement in tumor formation and proliferation [85–87]. Lgr5 induces the Wnt/ β-catenin pathway enhancing tumor formation and cancer cell proliferation [85]. The gradual increase in expression of Lgr5 we observed in our study was similar to that observed for cervical cancer [85]. Lgr5 expression increased as the disease progressed. Expression of CD44 also gradually increased as the disease progressed although the level of expression was lower compared to Lgr5. The present work showing a close association in expression between EMT transcriptional factors and stem cells markers is in agreement with studies indicating a link between EMT and acquisition of stem cell properties [8, 9]. In addition, studies have indicated that EMT facilitates generation of CSC traits for metastasis but also for self-renewal properties needed for initiating secondary tumors attributed to NANOG, OCT 4, SOX2 to name a few [90–92]. These gives credence to our observation that these markers can be used to detect gastric cancer metastasis and predict aggressive and fast progressing gastric cancers.

To date there are no published data showing the stage at which gastric cancer metastasizes. Using our mouse models of gastric cancer [42], we detected expression of EMT, stem cell markers and cytokeratins in our fast “progressors” gastric cancer model by 4 months but not in the slow “progressors” suggesting that these factors are involved in early events of tumorigenesis and therefore these factors may represent early indicators of disease dissemination and therefore metastasis. Our present work in mice suggests that dysplastic gastric epithelial cells start seeding themselves in other tissues including the bone marrow early during the disease progression to gastric cancer and before the emergence of gastric cancer in situ. In addition, our studies show an association between cytokeratins, EMTs, and CSCs with an aggressive form of gastric cancer. This study sets up a proof of concept that longitudinal monitoring of CTCs as an indicator of metastasis in gastric cancer is an achievable goal similar to the current management of breast cancer [20, 21, 28]. Therefore, findings from this study will lead to the development of early detection strategies for CTCs in patients with an aggressive form of gastric cancer so that appropriate treatment can be provided in a timely manner.

## Acknowledgements

This work is supported by the National Cancer Institute of the National Institute of Health under award R21CA210227. We wish to thank the UCSD School of Medicine Microscopy Core for access to the Keyence Immunofluorescence Microscope, which is supported by NINDS P30 Grant (NS047101).

## Abbreviations

*H. pylori*: (Helicobacter pylori)
WHO: (World Health Organization)
BMDCs: (Bone marrow derived cells)
*Myd88*^-/-^: (Myeloid differentiation primary response 88- deficient)
*H. felis*: *(Helicobacter felis)*
CTCs: (Circulating Tumor Cells)
DTCs: (Disseminating Tumor Cells)
EMT: (Epithelial to Mesenchymal Transition)
CK-8/-18/-19: (Cytokeratin 8/18/19)
WT: (Wild type)
c-Kit: (CD117)
MUC1: (Mucin 1)
ICC: (Immunocytochemistry)
qRT-PCR: (quantitative Real-time Polymerase chain reaction)
BHI: (Brain heart infusion)
FBS: (fetal bovine serum)
PBS: (phosphate buffered saline)
CD133: (Prominin 1)
GAPDH: (Glyceraldehyde 3-phosphate dehydrogenase)
MSC: (Mesenchymal Stem Cells)

## Author Contributions

All listed authors have made an impactful and substantial contribution to this work. All authors have approved the final manuscript for publication.

