## Supplementary Table 1 for "Early detection of biomarkers for circulating tumor cells in Bone marrow and Peripheral blood in a fast-progressing gastric cancer model"

**Supplementary Data**

**Supplementary Table 1: Primers used for RT-PCR analysis.**

| | **Sequence 5' – 3'** | **Reference** |
|---|---|---|
| **GAPDH** | F: TCAACAGCAACTCCCACTCTTCCA<br>R: ACCCTGTTGCTGTAGCCGTATTCA | **[1]** |
| **CD44** | F: CCACAGCCTCCTTTCAATAACC<br>R: GGAGTCTTCGCTTGGGGTA | **[2]** |
| **SOX9** | F: GGCAAGCTCTGGAGGCTG<br>R: CCTCCACGAAGGGTCTCTTCT | **[3]** |
| **Prominin1 (CD133)** | F: TCT GTT CAG CAT TTC CTC AC<br>R: TCA GTA TCG AGA CGG GTC | **In this study** |
| **SOX2** | F: AGGGTCTGCTACTGAGATGCTCTG<br>R: CAACCACTGGTTTTTCTGCCACCG | **[3]** |
| **OCT4** | F:TCTTTCCACCAGGCCCCCGGCTC<br>R:TGCGGGCGGACATGGGGAGATCC | **[3]** |
| **NANOG** | F: AGGGTCTGCTACTGAGATGCTCTG<br>R: CAACCACTGGTTTTTCTGCCACCG | **[3]** |
| **CK18** | F: ACTCCGCAAGGTGGTAGATG<br>R: GCCTCGATTTCTGTCTCCAG | **[4]** |
| **CK19** | F: ACCCTCCCGAGATTACAACC<br>R: CAAGGACTGTTCTGTCTCAA | **[5]** |
| **MUC1** | F: CCCTACCTACCACACTCACGGACG<br>R: GTGGTCACCACAGCTGGGTTGGTA | **[5]** |
| **Lgr5** | F: TGCCCATCACACTGTCACTGT<br>R: CACCCTGAGCAGCATCCTG | **[2]** |
| **Snail** | F: CTG GTG AGA AGC CAT TCT CC<br>R: GGA AGA TGC CAG CGA GGA TG | **In this study** |
| **Twist** | F: CTG GAC TCC AAG ATG GCA AG<br>R: CCA GAG TCT CTA GAC TGT CC | **In this study** |
| **ZEB** | F: CCA GTG AAG GTG ATC CAG CC<br>R: GAG GCC TCT TAC CTG TGT GC | **In this study** |

**Supplementary Figure Legends**

**S Fig 1: Bone marrow inmmunofluorescence of WT mice.** Representative images of immunofluorescent staining of bone marrow with CK8/18 (red), c-Kit (green) and nuclei stained with DAPI in wild type (WT) mice infected with *H. felis*, Images shown uninfected (a), 5 months post infection (b) and 6 months post infection (c).

**S. Fig 2: *Myd88*$^{-/-}$ mice infected with *H. felis* for 2 months.** Bone marrow was isolated from *Myd88*$^{-/-}$ mice , 2 months post infection and stained with CK8/18 (red), c-Kit (green) and nuclei stained with DAPI.
