## Supplementary figures and images for "Early detection of biomarkers for circulating tumor cells in Bone marrow and Peripheral blood in a fast-progressing gastric cancer model"

### S.Fig 1

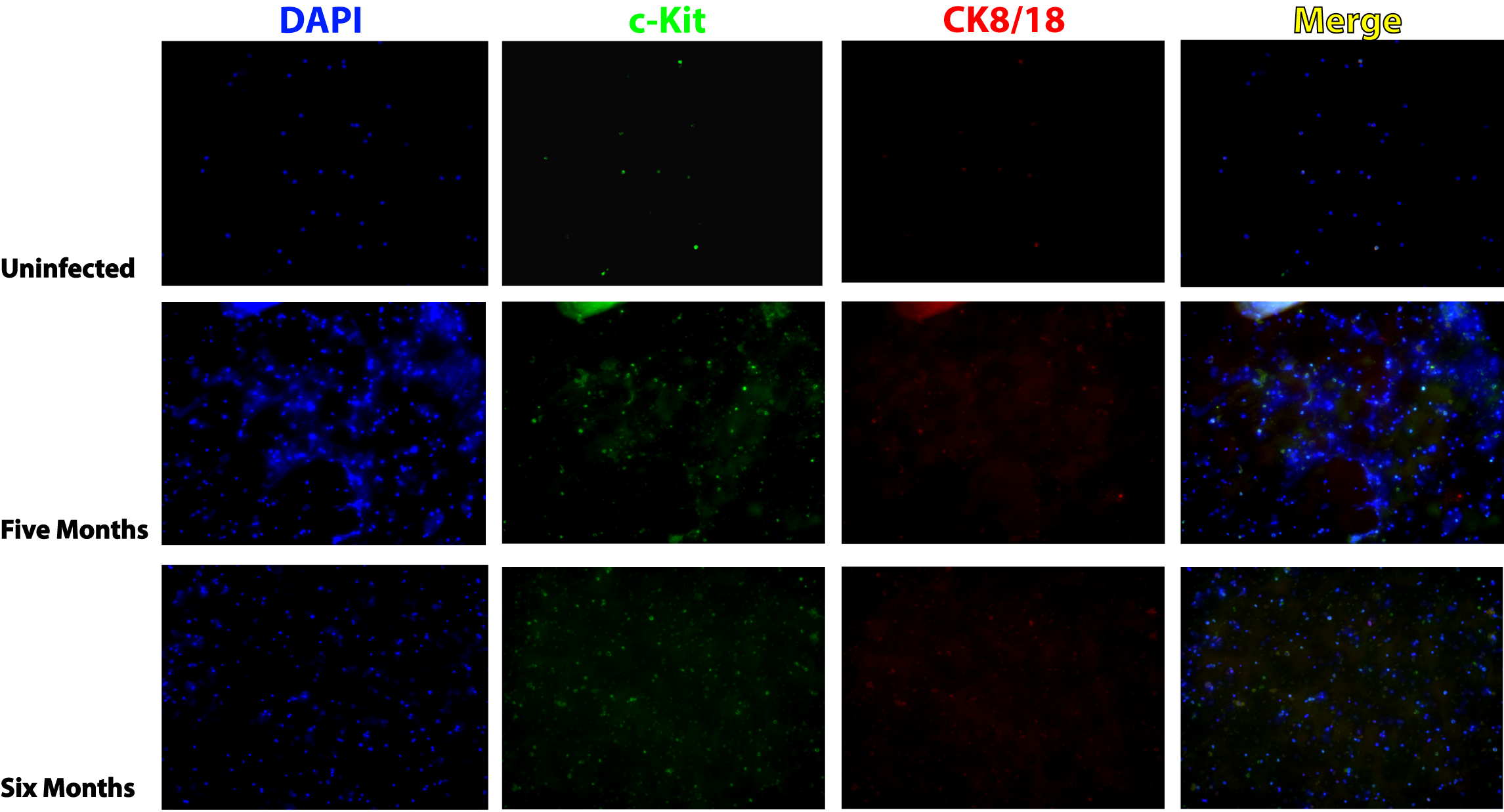

### S.Fig 2

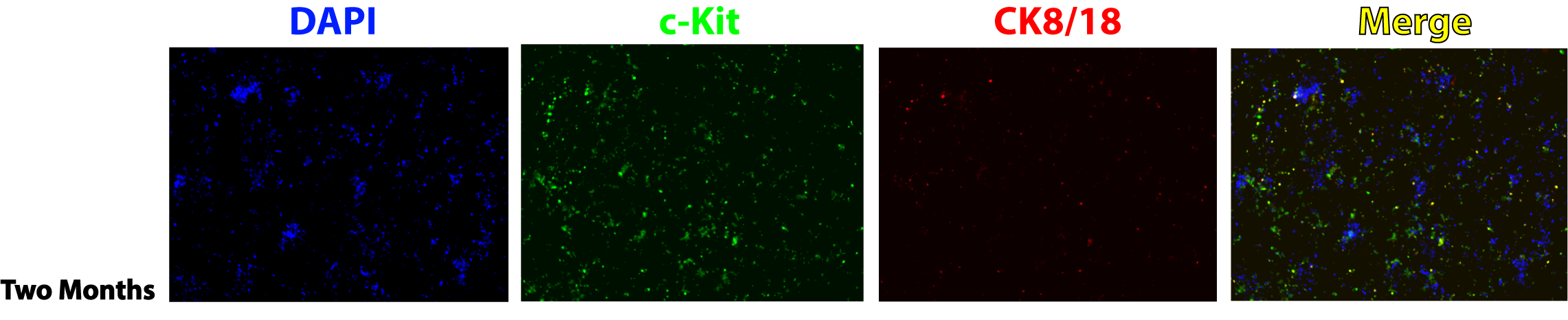
